# Expert programmers have fine-tuned cortical representations of source code

**DOI:** 10.1101/2020.01.28.923953

**Authors:** Yoshiharu Ikutani, Takatomi Kubo, Satoshi Nishida, Hideaki Hata, Kenichi Matsumoto, Kazushi Ikeda, Shinji Nishimoto

## Abstract

Expertise enables humans to achieve outstanding performance on domain-specific tasks, and programming is no exception. Many have shown that expert programmers exhibit remarkable differences from novices in behavioral performance, knowledge structure, and selective attention. However, the underlying differences in the brain are still unclear. We here address this issue by associating the cortical representation of source code with individual programming expertise using a data-driven decoding approach. This approach enabled us to identify seven brain regions, widely distributed in the frontal, parietal, and temporal cortices, that have a tight relationship with programming expertise. In these brain regions, functional categories of source code could be decoded from brain activity and the decoding accuracies were significantly correlated with individual behavioral performances on source-code categorization. Our results suggest that programming expertise is built up on fine-tuned cortical representations specialized for the domain of programming.

## 1. Introduction

Programming expertise is one of the most notable capabilities in the current computerized world. Since human software developers keep playing a central role in every software project and directly impact its success, this relatively new type of expertise is attracting increasing attention from modern industries (Li et al., 2015; Baltes and Diehl, 2018) and educations (Heintz et al., 2016; Papavlasopoulou et al., 2018). Moreover, huge productivity variations were repeatedly found even between programmers with the same level of experience (Boehm and Papaccio, 1988; DeMarco and Lister, 2013). Previous studies showed the psychological characteristics of expert programmers in their behaviors (Vessey, 1985; Koenemann and Robertson, 1991), knowledge structures (Fix et al., 1993; Von Mayrhauser and Vans, 1995), and eye movements (Uwano et al., 2006; Busjahn et al., 2015). Although these studies clearly illustrate the behavioral specificity of expert programmers, it remains unclear what neural bases differentiate expert programmers from novices.

Recent studies have investigated the brain activity of programmers using fMRI to examine their cognitive mechanisms. Siegmund *et al*. contrasted brain activity during program output estimations against syntax error searches and showed that the processes of program output estimations activated left-lateralized brain regions (Siegmund et al., 2014, 2017). Several studies have tried to investigate neural correlates of subject-wise programming expertise but failed to find a systematic trend (Peitek et al., 2018a). Although an exploratory study argued the correlation between activity pattern discriminability and students’ GPA score (Floyd et al., 2017), the assumed relationship of GPA scores to programming expertise was ambiguous and not empirically validated. Further, the main limitation of these prior studies is the use of a single homogeneous subject group that only covered a small range of programming expertise. Recruitment of more diverse subjects in terms of their programming expertise may enable us to elucidate the potential differences of brain functions related to the expertise (Bilalić, 2017).

Here we aim to identify the neural bases of programming expertise that contribute outstanding performances of expert programmers and provide a clue to describe how the brain accommodates such behavioral superiority in programming. To do this, we defined two fundamental factors in our experiment: An objective indicator of programming expertise and a laboratory task that efficiently exhibits experts’ superior performances under the general constraints of fMRI experiments. For the first factor, we adopted programmers’ ratings in competitive programming contests (AtCoder), which are objectively determined by the relative positions of their actual performances among thousands of programmers (Wasik et al., 2018). We recruited top- and middle-rated programmers as well as novice controls to cover a wide range of programming expertise in our fMRI experiment. For the second factor, we developed the program categorization task and confirmed that behavioral performances of this task were significantly correlated with the programming expertise indicator. This confirmation allows us to expect the tight association between individual programming expertise and the brain activity patterns recorded by fMRI while subjects performed this laboratory task.

To examine the brain activity patterns underlying expert programmers’ behavioral superiority, we employ a decoding framework that learns the relationship between multivoxel activity patterns in the brain and functional categories of source code. Our hypothesis is that higher programming expertise relates to specific multi-voxel pattern representations, potentially influenced by their domain-specific knowledge and training experiences. This framework is motivated by prior studies that contrasted multi-voxel activity patterns of experts against novices and demonstrated that domain-specific expertise generally associates with representational changes in the brain (Bilalić et al., 2016; de Borst et al., 2016; Martens et al., 2018; Gomez et al., 2019). In the present study we adopt whole-brain searchlight analysis (Kriegeskorte et al., 2006) to explore the potential loci of programming expertise in a data-driven manner.

Here we demonstrate that functional categories of program source code can be decoded from programmers’ brain activity and decoding accuracies in seven distinct brain regions are significantly correlated with individual behavioral performances that reflect programming expertise. Furthermore, we show that decoding accuracies of subordinate-level categories on two brain regions are significantly correlated with individual behavioral performance, even though such discriminations are not explicitly required by the tasks. These results suggest that expert programmers’ outstanding performances depend on fine-tuned cortical representations of source code and such cortical representation refinements might be related to the acquisition of advanced-level programming expertise.

## 2. Materials and methods

### 2.1. Subjects

Thirty healthy subjects (two females, aged between 20 and 24 years) participated in the experiment; see Table.1 for the detailed demographic information. Ten subjects were top 20% rankers (*Expert*) and another ten were 21-50% rankers (*Middle*) in AtCoder (https://atcoder.jp/), based on the ranking at July 1 2017. Ten control subjects (*Novice*) with four years or less programming experience and no experience on competitive programming were also included. All were right-handed (assessed by the Edinburgh Handedness Inventory (Oldfield, 1971), laterality quotient = 83.6 ± 24.0, ranged between +5.9 and +100) and understood basic Java grammars with at least half of year experience on Java programming. The averaged AtCoder rates (1,967 in *Expert* and 894 in *Middle*) were equivalent to the top 6.5% and 34.1% positions among 7,671 registered players, respectively. Seven additional subjects were scanned but not included in the analysis because one showed neurological abnormality in MRI images, three retired the experiment without full completion, three showed strongly-biased behavioral responses judged when the behavioral performance of one or more choices did not reach chance-level in the training experiments, signaling the strong response bias sticking to a specific choice. This study was approved by the Ethics Committees of NICT and NAIST and subjects gave written informed consent for participation. The sample size was chosen to match previous fMRI studies on human expertise with similar behavioral protocols (Amalric and Dehaene, 2016; Bilalić et al., 2016; de Borst et al., 2016).

### 2.2. Stimuli

For this study 72 code snippets written in Java were collected from an open codeset provided by AIZU ONLINE JUDGE (http://judge.u-aizu.ac.jp/onlinejudge/); an online judge system where lots of programming problems are listed and everyone can submit their own source code to answer those online. We selected four functional categories (*category*) and eleven subordinate concrete algorithms (*subcategory*) based on two popular textbooks about computer algorithms (Cormen et al., 2009; Sedgewick and Wayne, 2011); see Fig.1a and Supplementary Table 1 and 2 for the detailed descriptions. We first searched in the open codeset for Java code snippets implementing one of the selected algorithms and found 1251 candidates. The reasons why we focused on Java in this study were because the language has been one of the most famous programming languages and prior fMRI studies on programmers also used Java code snippets as experimental stimuli (Siegmund et al., 2014, 2017; Peitek et al., 2018a). To meet the screen size constraint in the MRI scanner, we excluded code snippets with lines of code (LOC) more than 30 and characters per line (CPL) more than 120. From all remaining snippets, we created a set of 72 code snippets with minimum deviations of LOC and CPL to minimize visual variation as experimental stimuli; the mean and standard deviation of LOC and CPL were 26.4 ± 2.4 and 59.3 ± 17.1, respectively. In the codeset, 18 snippets each belonged to one of the *category* classes and six snippets each belonged to one of the *subcategory* classes except for “linear search” class with twelve snippets (see Supplementary Table 3 for statics of the codeset). The indentation styles of all code snippets were normalized by replacing a tab-space with two white-spaces and all user-defined functions were renamed to neutral like “function1” because some of them indicated their algorithms explicitly (see Supplementary Figure 1 for example snippets used in the experiment). We verified all code snippets had no syntax error and run correctly without run-time error.

**Figure 1:**
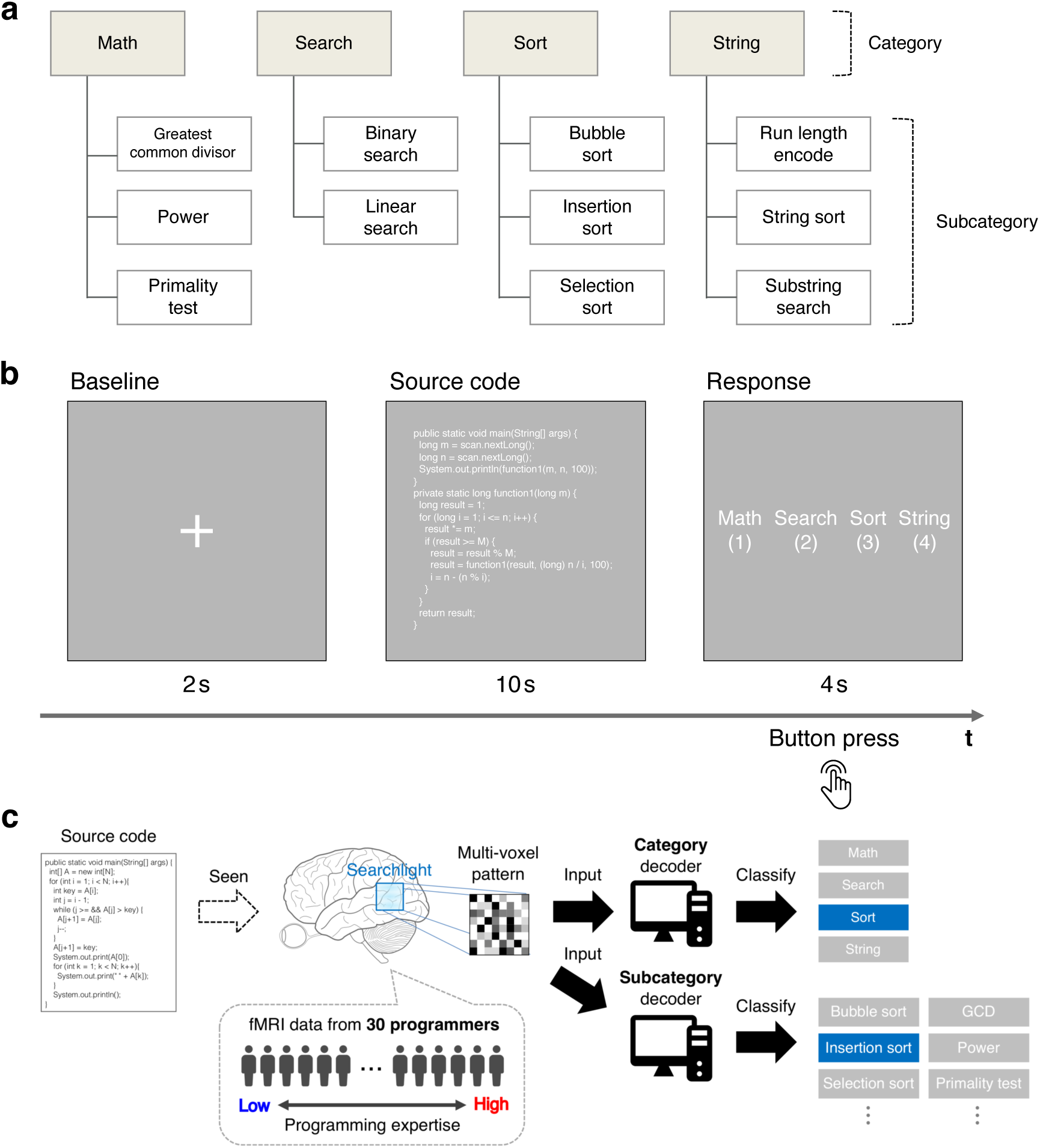
Experimental design. (a) Hierarchy of categories used in this study. Category and Subcategory represent abstract functionality and concrete algorithms, respectively, based on two popular textbooks of programming. Every code snippet used in this study belonged to one subcategory class and its corresponding category class. (b) Program categorization task. After a fixation-cross presentation for two seconds, a Java code snippet was displayed for ten seconds in white text without any syntax highlight. Then, subjects responded the category of given code snippet by pressing a button. (c) Overview of the decoding framework. MRI data was collected from 30 subjects with different levels of programming expertise while they performed the program categorization task. Whole-brain searchlight analysis (Kriegeskorte et al., 2006) was employed to explore the potential loci of programming expertise. For each searchlight location, a linear-kernel SVM classifier (decoder) was trained on multi-voxel patterns to classify *category* or *subcategory* of given Java code snippets.

**Table 1:** Demographic information of recruited subjects. Numerics from 4th (Age) to last columns denote ‘MEAN ± SD’. Abbreviations: PE, programming experience; JE, Java experience; CPE, competitive programming experience [Year]. Significant differences were observed between PE of Expert - Novice, Middle - Novice; CPE of Expert - Middle (two-sample t-test, p < 0.05 FDR-corrected).

|  | N | Sex (M/F) | Age | AtCoder rate | PE | JE | CPE |
| --- | --- | --- | --- | --- | --- | --- | --- |
| Expert | 10 | 10 / 0 | 22.6 ± 1.1 | 1969 ± 467 | 6.9 ± 2.8 | 2.8 ± 2.4 | 4.1 ± 2.6 |
| Middle | 10 | 9 / 1 | 22.5 ± 0.8 | 894 ± 175 | 4.8 ± 1.7 | 1.1 ± 0.8 | 1.3 ± 0.8 |
| Novice | 10 | 9 / 1 | 21.7 ± 1.2 | NA | 2.8 ± 0.6 | 1.4 ± 1.0 | NA |

### 2.3. Experimental design

The fMRI experiment consisted of six separate runs (9 min 52 sec for each run). Each run contained 36 trials of the program categorization task (Fig.1b) plus one dummy trial to avoid undesirable effects of MRI signal instability. We used 72 code snippets as stimuli and each snippet was presented three times through the whole experiment (216 trials in total), but the same snippet appeared only once in a run. We employed PsychoPy (Peirce, 2007) (version 1.85.1) to display the code snippets in white text and gray background without any syntax highlighting to minimize visual variations. In each trial of program categorization tasks, a Java code snippet was displayed for ten seconds after a fixation-cross presentation for two seconds. Subjects then responded via pressing buttons placed under the right hand to indicate which *category* class was most plausible for the code snippet and all response data were automatically collected for the calculation of individual behavioral performance. To clarify classification criteria, a brief explanation about each *category* class was provided before the experiment started (see Supplementary Table 1). The presentation order of the code snippets was randomized under balancing the number of exemplars for each *category* class across runs. The corresponding buttons for each answer choice were also randomized across trials to avoid linking a specific answer choice with a specific finger movement. Subjects were allowed to take a break between runs and to quit the fMRI experiment at any time.

To mitigate potential noises caused by task unfamiliarity, every subject conducted a training experiment within ten days before the fMRI experiment. The training experiment consisted of three separate runs with the same settings as the fMRI experiment. A different set of 72 Java code snippets implementing the same algorithms was used as stimuli; each snippet was presented one or two times but the same snippet appeared only once in a run. In addition, all subjects took a post-MRI experiment within ten days after the fMRI experiments for assessment of individual ability to *subcategory* categorizations. Before the post-MRI experiments started, we revealed the existence of *subcategory* and assessed whether the subjects recognized subcategory classes during the fMRI experiment using a questionnaire. The post-MRI experiment consisted of two separate runs using the same codeset as the fMRI experiment. Subjects were provided brief descriptions about each *subcategory* class (see Supplementary Table 2) and classified the given code snippet from two or three choices of *subcategory* classes according to its superordinate *category*, e.g. ‘bubble sort’, ‘insertion sort’, ‘selection sort’ were displayed when the snippet in ‘sort’ category was presented. The training and post-MRI experiments were performed outside of the MRI scanner. For all experiments, we calculated behavioral performance as a ratio of correct-answer-trials in all-trials; unanswered trials were regarded as ‘incorrect’ for this calculation. Chance-level behavioral performance was 25% in the training and fMRI experiments and 37.25% in the post-MRI experiment adjusted for imbalanced numbers of answer choices.

### 2.4. MRI data acquisition

MRI data were collected using 3-Tesla Siemens MAGNETOM Prisma scanner with a 64-channel head coil located at CiNet. T2^*\**^-weighted multiband gradient echo-EPI sequences were performed to acquire functional images covering the entire brain (repetition time (TR) = 2000 ms, echo time (TE) = 30 ms, flip angle = 75°, field of view (FOV) = 192 × 192 mm, slice thickness = 2 mm, slice gap = 0 mm, voxel size = 2 × 2 × 2.01 mm, multi-band factor = 3). A T1-weighted magnetization-prepared rapid acquisition with gradient-echo sequence was also performed to acquire fine-structural images of the entire head (TR = 2530 ms, TE = 3.26 ms, flip angle = 9°, FOV = 256 × 256 mm, slice thickness = 1 mm, slice gap = 0 mm, voxel size = 1 × 1 × 1 mm).

### 2.5. MRI data preprocessing

We used the Statistical Parametric Mapping toolbox (SPM12, http://www.fil.ion.ucl.ac.uk/spm/) for preprocessing. The first eight scans in dummy trials for each run were discarded to avoid MRI signal instability. The functional scans were aligned to the first volume in the fourth run to remove movement artifacts. They were then slice-time corrected and co-registered to the whole-head T1 structural image. Both anatomical and functional images were spatially normalized into the standard Montreal Neurological Institute 152-brain average template space and resampled to a voxel size of 2 × 2 × 2 mm. MRI signals at each voxel were high-pass–filtered with a cutoff period of 128 seconds to remove low-frequency drifts. A thick gray matter mask was obtained from the normalized anatomical images of all subjects to select the voxels within neuronal tissue using the SPM Masking Toolbox (Ridgway et al., 2009). For each subject independently, we then fitted a general linear model (GLM) to estimate voxel-level parameters (*β*) linking recorded MRI signals and conditions of source code presentations in each trial. The fixation and response phases in each trial were not explicitly modeled. The model also included motion realignment parameters to regress-out signal variations due to head motion. Finally, 216 beta estimate maps (36 trials × 6 runs) per subject were yielded and used as input for the following multivariate pattern analysis.

### 2.6. Multi-voxel pattern analysis

We used whole-brain searchlight analysis (Kriegeskorte et al., 2006) to examine where significant decoding accuracies exist using the Decoding Toolbox (Hebart et al., 2015) (version 3.99) and LIBSVM (Chang and Lin, 2011) (version 3.17). A four-voxel-radius sphered searchlight, covering 251 voxels at once, was systematically shifted throughout the brain and decoding accuracy was quantified on each search-light location. A linear-kernel SVM classifier was trained and evaluated using a leave-one-run-out cross-validation procedure, which iteratively treated data in a single run for test and others for training. In each fold, training data was first scaled to zero-mean and unit variance by z-transform and test data was scaled using the estimated scaling parameters. We then applied outlier reduction using [-3, +3] as cut-off values and all scaled signals larger than the upper cut-off or smaller than the lower cut-off were set to the closest value of these limits. The SVM classifier was trained with three cost parameter candidates [0.1, 1, 10] and the best parameter was chosen by grid search in nested cross-validations. We here adopted a relatively small set of parameter candidates due to the constraint of the high computational load of searchlight analysis. Finally, the trained classifier predicted *category* or *subcategory* of seen source code from the leave-out test data and decoding accuracy was calculated as a ratio of correct-classifications out of all-classifications. Note that corrected misclassification cost weights were used in *subcategory* decoding to compensate for the imbalanced number of exemplars across *subcategory* classes.

The training and evaluation procedures were performed independently for each subject and a whole-brain decoding accuracy map was obtained per subject. We then conducted second-level analyses to examine the significance of decoding accuracies and correlations between individual decoding and behavioral performances. For this purpose, the decoding accuracy maps were spatially smoothed using a Gaussian kernel of 6 mm full-width at half maximum (FWHM) and submitted to random effects analysis as implemented in SPM12. The analysis tested the significance of group-level decoding accuracy and Pearson’s correlation coefficient between individual decoding accuracies and behavioral performances. A relatively strict statistical threshold of voxel-level p < 0.05 FWE-corrected was used for decoding accuracy tests and a standard threshold of voxel-level p < 0.001 uncorrected and cluster-level p < 0.05 FWE-corrected was used for correlation tests. The chance-level accuracy (25% in *category* decoding and 9.72% in *subcategory* decoding; adjusted for imbalanced numbers of exemplar) and zero correlation were adopted as null hypotheses.

### 2.7. Data and code availability

The experimental data and code used in the present study are available from our repository: https://github.com/Yoshiharu-Ikutani/DecodingCodeFromTheBrain.

## 3. Results

### 3.1. Behavioral data

We evaluated the relationship between the programming expertise indicator and behavioral performance on the program categorization task. A significant correlation was observed between AtCoder rate (M = 954.3, SD = 864.6) and behavioral performance in the fMRI experiments (M = 76.0, SD = 13.5 [%]), r = 0.593, p = 0.0059, n = 20 (Fig.2a). The correlation was kept if we included behavioral performances of non-rate-holders (i.e. novices) as zero-rated subjects; r = 0.722, p = 0.000007, n = 30. We additionally found a positive correlation between AtCoder rate and behavioral performance on subcategory categorization in the post-MRI experiments (M = 65.9, SD = 17.0 [%]), r = 0.688, p = 0.0008, n = 20 (Fig.2b). The significant correlation was also kept if we included non-rate-holder subjects; r = 0.735, p = 0.000004, n = 30. This result was consistent with the self-reporting data indicating that subjects with higher programming expertise recognized more *subcategory* classes during the fMRI experiments (Supplementary Table 4). From all behavioral data, we certainly concluded that behavioral performances on the program categorization task significantly correlated with programming expertise. The behavioral evidence allowed us to expect that individual programming expertise was reflected in the brain activity patterns measured using fMRI while subjects performed this laboratory task.

**Figure 2:**
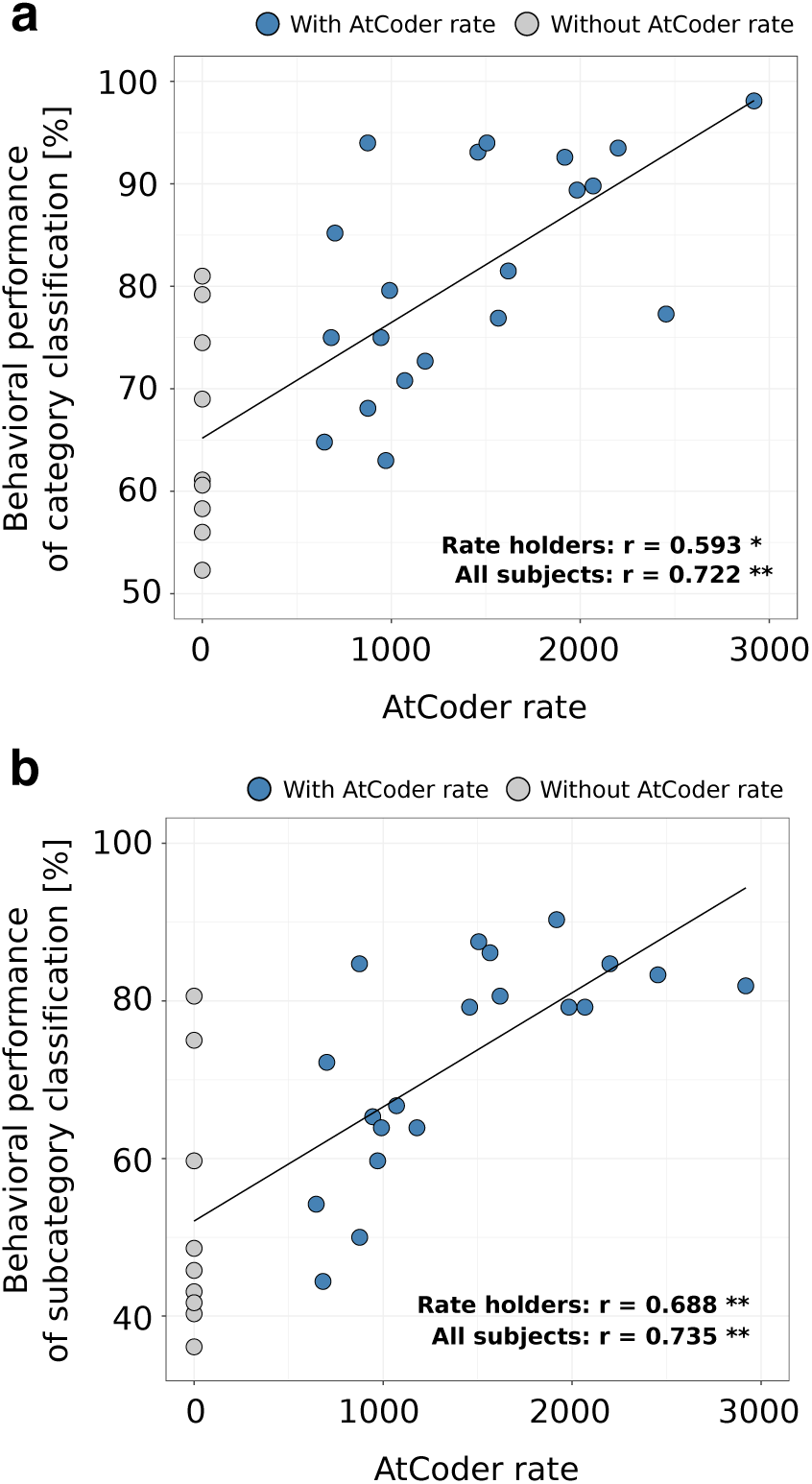
Correlations between behavioral performance and programming expertise indicator. (a) Scatter plot of behavioral performances of category classifications against the values of expertise indicator (i.e. AtCoder rate). (b) Scatter plot of behavioral performances of subcategory classifications against the values of expertise indicator. Each dot represents an individual subject. One-sample t-tests were used to check significance of the correlation coefficients (r) between the expertise indicator and behavioral performances; *, p < 0.05 and **, p < 0.005. The solid lines indicate a fitted regression line estimated from all subject data.

### 3.2. Expertise-related mutli-voxel patterns

We first examined where we could decode the functional categories of source code from programmers’ brain activity. Fig.3 visualizes the searchlight centers that showed significantly high decoding accuracy than chance estimated from all subject data using a relatively strict whole-brain statistical threshold (voxel-level p < 0.05 FWE-corrected). The figure shows that significant decoding accuracies were observed in the broad areas of bilateral occipital cortices, parietal cortices, posterior and ventral temporal cortices, as well as the bilateral frontal cortices around inferior frontal gyri. Given the result, we confirmed that functional categories of source code were represented in the widely distributed brain areas and the cortical representations of each *category* class were linearly separable by a simple SVM classifier.

**Figure 3:**
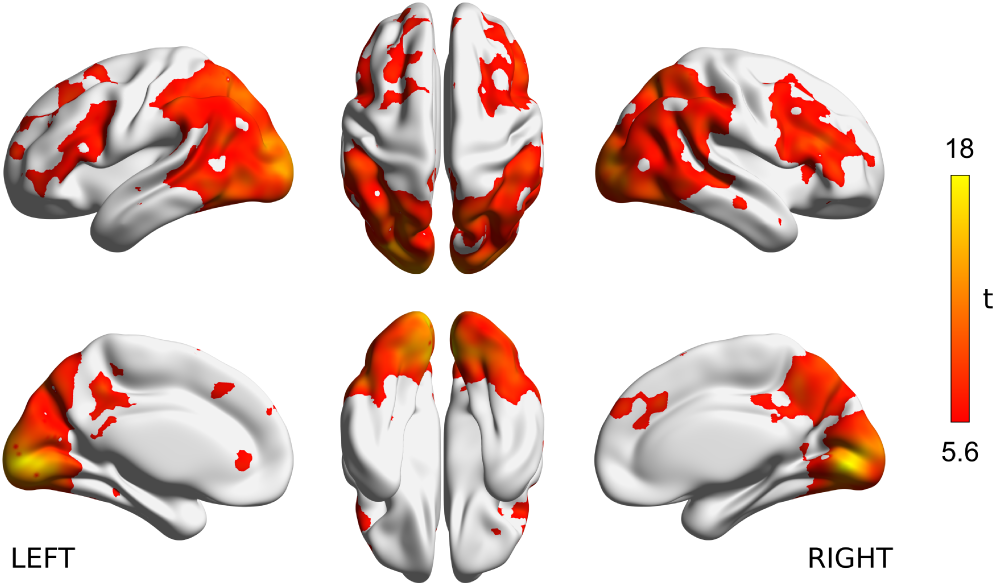
Decoding accuracy for functional category of source code. (a) Significant searchlight locations estimated from all subject data (N = 30). Heat colored voxels denote the centers of searchlights with significant decoding accuracy (voxel-level p < 0.05, FWE corrected). See Supplementary Figure 2a for the distribution of voxel-level peak decoding accuracies. The brain surface visualizations were performed using BrainNet viewer, version 1.61 (Xia et al., 2013).

To associate the cortical representation of source code with individual programming expertise, we investigated a linear correlation between behavioral performances and decoding accuracies for each searchlight location. Fig.4a visualizes the searchlight centers that showed significantly high correlation coefficients using thresholds of voxel-level p < 0.001 uncorrected and cluster-level p < 0.05 FWE-corrected. We observed significant correlations in the areas of bilateral inferior frontal gyri pars triangularis (IFG Tri), right superior frontal gyrus (SFG), left inferior parietal lobule (IPL), left middle and inferior temporal gyrus (MTG / IT); see the slice-width visualization shown as Fig.4b and Supplementary Table 5 for the list of significant clusters. In this correlation analysis, the right IFG Tri showed the highest peak correlation coefficient (r = 0.79, p < 10^−6^, Fig.4c). These results provided evidence that cortical representations in the distinct brain areas mainly located in frontal, parietal, and temporal cortices were significantly associated with experts’ outstanding performances on the program categorization task. In contrast, cortical representations in the bilateral occipital cortices including early visual areas did not show a significant correlation to individual behavioral performances, while significant decoding accuracies were broadly observed in the cortices shown as Fig.3a.

**Figure 4:**
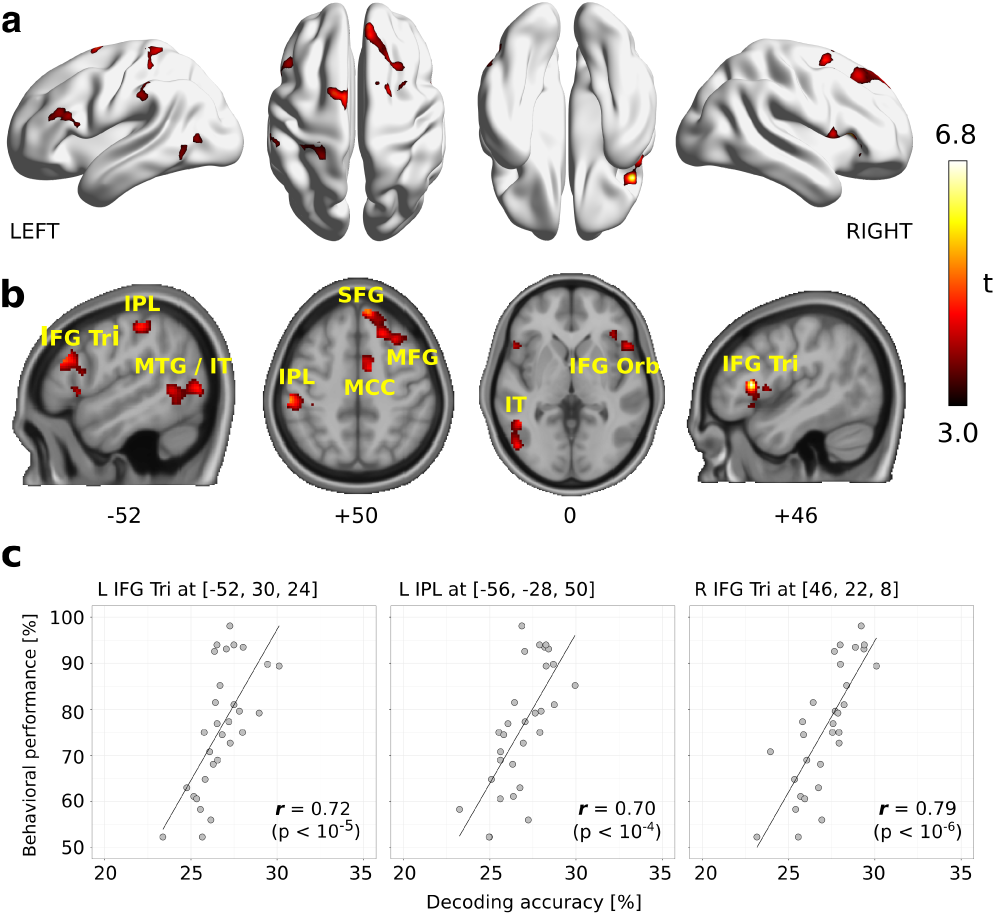
Searchlight-based correlation analysis between behavioral performances and decoding accuracies. (a) Locations of searchlight showing significant correlations. Significance was determined by a threshold of voxel-level p < 0.001 and cluster-level p < 0.05, FWE corrected for the whole brain. (b) Slice-wise visualizations of the significant clusters using bsp- mview (http://www.bobspunt.com/software/bspmview). (c) Scatter plots of peak correlations between decoding accuracies and behavioral performances. Each dot represents an individual subject data. Correlation coefficients (r) and uncorrected p values are shown in bottom-right of each plot. See Supplementary Table 5 and Supplementary Figure 3 for all significant clusters and peak correlations. Abbreviations: SMG, Supramarginal gyrus; IPL, Inferior parietal lobule; MTG, Middle temporal gyrus; IT, Inferior temporal gyrus; SFG, Superior frontal gyrus; MFG, middle frontal gyrus; IFG Tri, Inferior frontal gyrus pars triangularis; IFG Orb, Inferior frontal gyrus pars orbitalis; MCC, medial cingulate cortex.

Previous two analyses separately showed where significant decoding accuracies exist and whether the decoding accuracies significantly correlate with behavioral performances. To achieve more validated evidence for the cortical representations associated with programming expertise, we integrated these two analyses and identified searchlight centers that had sufficient information to represent functional categories of source code and their decoding accuracies significantly correlated with individual behavioral performance. As a result, we found 1,205 searchlight centers (equal to 0.79%) that survived from both statistical thresholds of decoding accuracy and correlation to behavioral performances; shown as red-colored dots in Fig.5a. The survived searchlight centers were mainly observed in the bilateral IFG Tri, left IPL, left supramarginal gyrus (SMG), left MTG/IT, and right middle frontal gyrus (MFG) as shown in Fig.5b. The complementary *sensitivity* analysis (Etzel et al., 2013) using a five-voxel-radius searchlight showed the almost same tendency, indicating that the results were not limited to a specific searchlight radius parameter (see Supplementary Figure 4 and Supplementary Table 6). Since we have demonstrated that individual behavioral performances were significantly correlated with the expertise indicator in competitive programming contests (Fig.2a), this result revealed a tight association between high-level programming expertise and the improvement of decoding accuracy in these seven brain regions.

**Figure 5:**
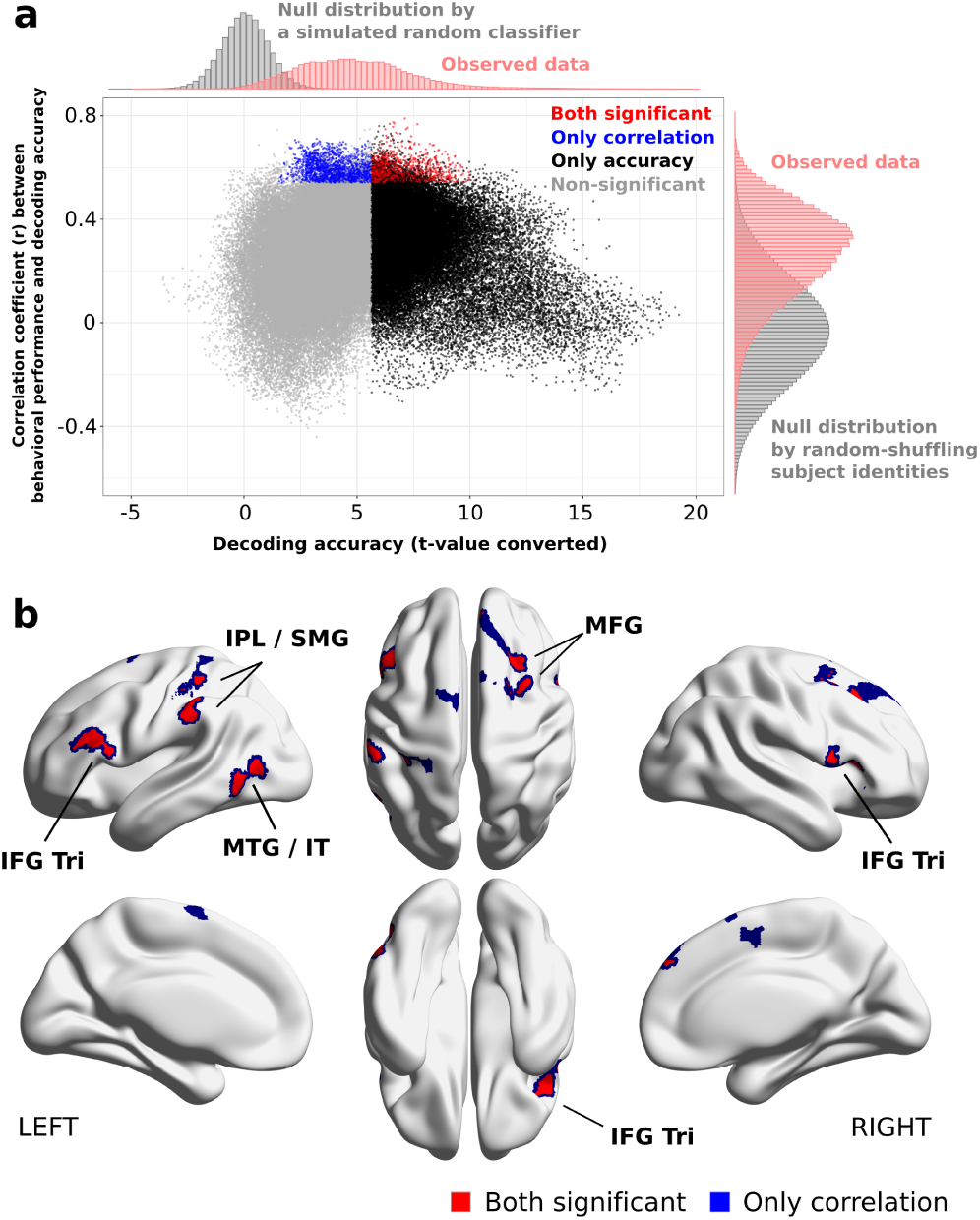
Identifying searchlight centers that showed both significant decoding accuracy and significant correlation to individual behavioral performances. (a) Scatter plot of searchlight results. X-axis shows t-values calculated from all subjects’ decoding accuracies on each searchlight locations. Y-axis indicates correlation coefficients between decoding accuracies and behavioral performances. Red-colored dots denote the searchlights showing both significant decoding accuracy and correlation, while blue and black denote those only showed significant decoding accuracy or correlations. Non-significant searchlights were colored in gray. The observed distributions of decoding accuracies and correlations are respectively shown on top- and right-sides of the figure accompanied with null distributions calculated by randomized simulations. (b) Locations of searchlight centers that showed both significant decoding accuracy and significant correlations to individual behavioral performances.

### 3.3. Representations of subcategory information

We next investigated where we could decode the *subcategory* of source code from programmers’ brain activity to examine finer-level cortical representations. In our experiment, subjects responded ‘sort’ when he/she has been presented with the code snippets implementing one of three different sorting algorithms; i.e. bubble, insertion, and selection sorts (Fig.1a). This cognitive process could be considered as a generalization process that incorporates different but similar algorithms (*subcategory*) into a more general functionality class (*category*). Additionally, several psychologists indicated that experts specifically show high performances in subordinate-level categorizations as well as basic-level categorizations (Tanaka and Taylor, 1991). In fact, we have observed that the ability to differentiate *subcategory* classes significantly correlated to the programming expertise indicator in competitive programming (Fig.2b). This evidence implies that programmers’ brain activity patterns may automatically respond to the detailed functional difference of source code. The decoding accuracy of *subcategory* may be correlated with programming expertise, even though they classified only *category* classes, not *subcategory*, of given code snippets and the existence of *subcategory* classes had never been revealed until the end of fMRI experiment.

We employed searchlight analysis with the same setting as used in the previous analysis to reveal the spatial distribution of significant *subcategory* decoding accuracies and significant correlations to behavioral performances. Fig.6 illustrates the searchlight centers that showed significantly high *subcategory* decoding accuracy than chance (9.72%; corrected for imbalanced exemplars) using a threshold of voxel-level p < 0.05 FWE-corrected. Linear correlation between *subcategory* decoding accuracies and individual behavioral performances was then assessed using thresholds of voxel-level p < 0.001 uncorrected and cluster-level p < 0.05 FWE-corrected (Fig.7). As a result, only a cluster on the left SMG and superior temporal gyrus (STG) showed a significant correlation; the peak correlation coefficient was observed in the left STG (r = 0.72, p < 10^−5^; Fig.7c). Finally, we integrated the results from decoding and correlation analysis of *subcategory* and confirmed that 120 searchlight centers (equal to 0.08%) on the left SMG and STG survived from both statistical thresholds of decoding accuracy and correlation to behavioral performances; shown as red-colored dots in Fig.8a. The complementary *sensitivity* analysis using a five-voxel-radius searchlight indicated that these results were consistently observed across the two searchlight radius parameters (see Supplementary Figure 5 and Supplementary Table 7). These results suggest that cortical representations of fine functional categories on the left SMG and STG may play an important role in achieving advanced-level programming expertise, even though the representations are not explicitly required by the tasks.

**Figure 6:**
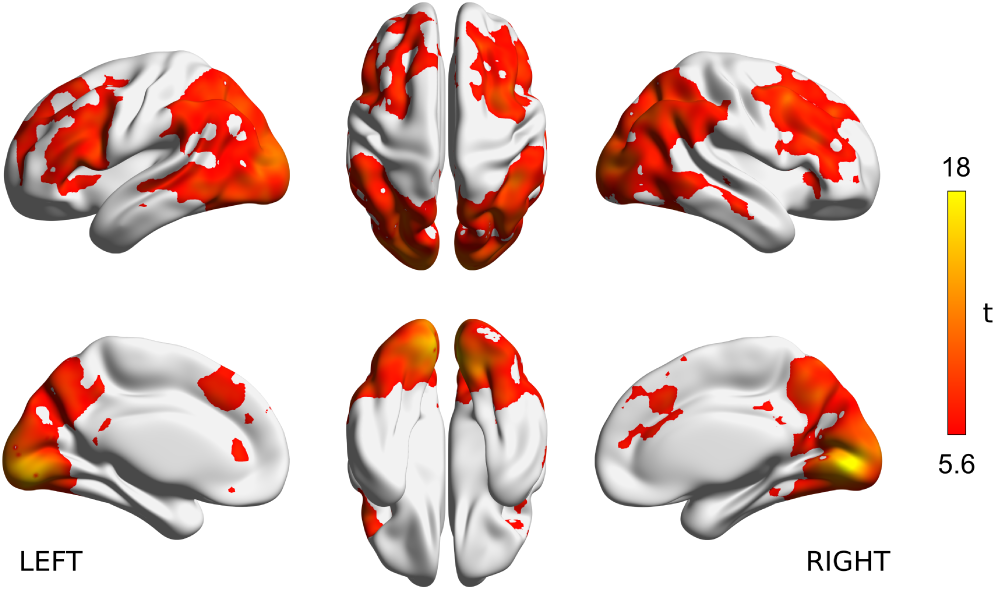
Decoding accuracy for subcategory of source code. (a) Searchlight locations showing significant subcategory decoding accuracy than chance estimated from all subject data (N = 30). Heat colored voxels denote the centers of searchlights with significant subcategory decoding accuracy (voxel-level p < 0.05, FWE corrected). See Supplementary Figure 2b for the distribution of voxel-level peak subcategory decoding accuracies.

**Figure 7:**
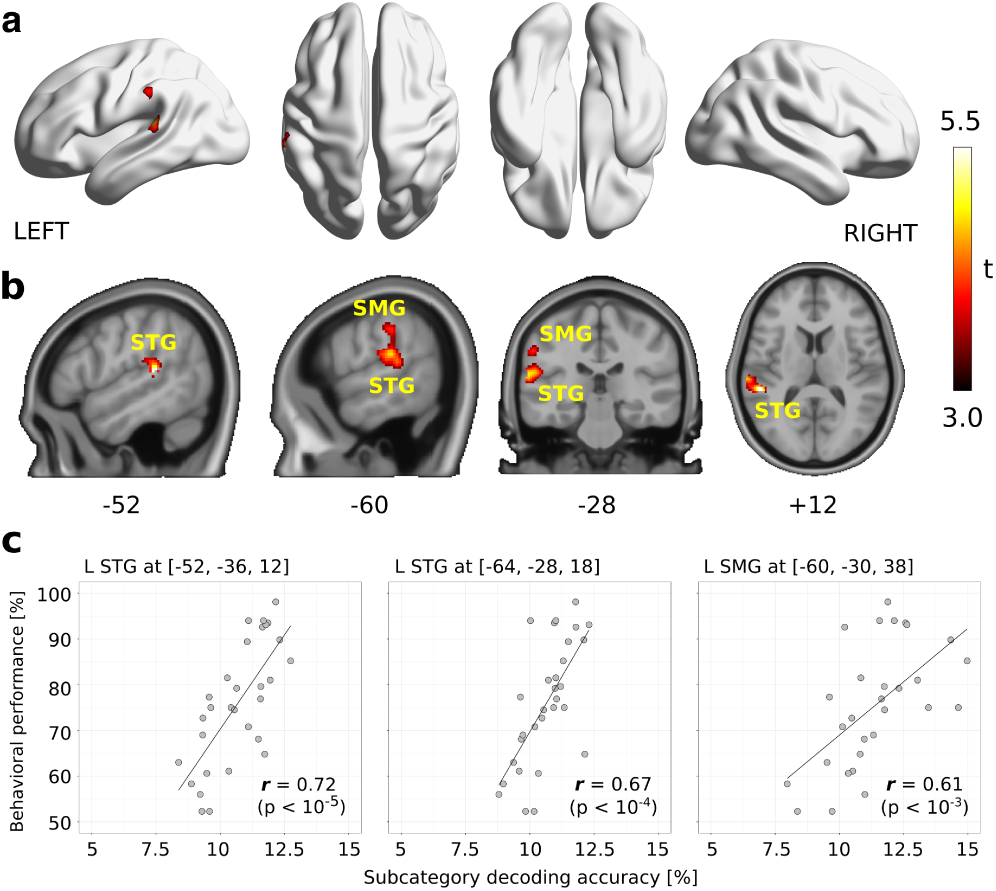
Searchlight-based correlation analysis between behavioral performances and subcategory decoding accuracies. (a) Locations of searchlight showing significant correlations. Significance was determined by a threshold of voxel-level p < 0.001 and cluster-level p < 0.05, FWE corrected for the whole brain. (b) Slice-wise visualizations of the significant clusters. (c) Scatter plots of peak correlations between decoding accuracies and behavioral performances. Each dot represents an individual subject data. Correlation coefficients (r) and uncorrected p values are shown in bottom-right of each plot. Only one cluster (extent = 501 voxels) had significant correlation in this analysis and three peak correlations in the cluster were shown here. Abbreviations: STG, Superior temporal gyrus.

**Figure 8:**
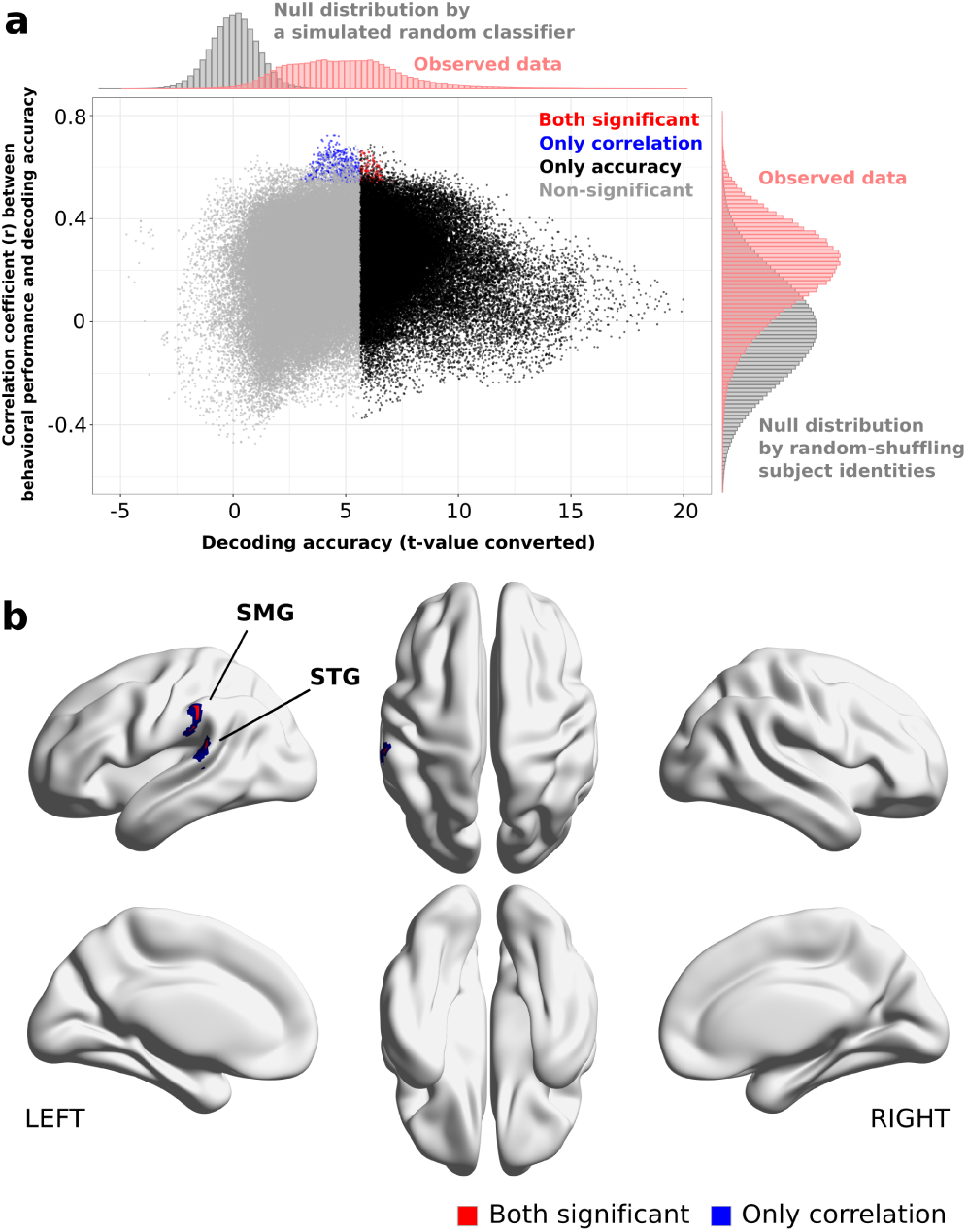
Identifying searchlight centers that showed both significant subcategory decoding accuracy and significant correlation to individual behavioral performances. (a) Scatter plot of searchlight results. X-axis shows t-values calculated from all subjects’ decoding accuracies on each searchlight locations. Y-axis indicates correlation coefficients between subcategory decoding accuracies and behavioral performances. Red-colored dots denote the searchlights showing both significant decoding accuracy and correlation, while blue and black denote those only showed significant decoding accuracy or correlations. Non-significant searchlights were colored in gray. The observed distributions of subcategory decoding accuracies and correlations are respectively shown on top- and right-sides of the figure accompanied with null distributions calculated by randomized simulations. (b) Locations of searchlight centers that showed both significant subcategory decoding accuracy and significant correlations to individual behavioral performances.

## 4. Discussion

We have shown that functional categories of source code can be decoded from programmers’ brain activity measured using fMRI. Decoding accuracies on the bilateral IFG Tri, left IPL, left SMG, left MTG, left IT, and right MFG were significantly correlated with individual behavioral performances on the program categorization task. Furthermore, decoding accuracies of *subcategory* on the left SMG and STG were also strongly correlated with the behavioral performances while the subordinate-level representations were not directly induced by the performing tasks. Since we have demonstrated the behavioral performances were correlated with the expertise indicator in competitive programming contests, our results revealed a tight association between advanced-level programming expertise and domain-specific cortical representations in these brain areas widely distributed in the frontal, parietal, and temporal cortices.

Previous fMRI studies on programmers have aimed at characterizing how programming-related activities, such as program comprehension and bug detection, take place in the brain (Siegmund et al., 2014, 2017; Floyd et al., 2017; Castelhano et al., 2019; Peitek et al., 2018a,b). Exceptionally, an exploratory study reported that BOLD signal discriminability between code and text comprehensions was negatively correlated with participants’ GPA scores in a university (Floyd et al., 2017). However, the relationship between GPA scores and programming expertise was ambiguous and the observed correlation was relatively small (r = -0.44, p = 0.016, n = 29). Our aim in the present study was substantially different: We sought the neural bases of programming expertise that contribute expert programmers’ outstanding performances. To address the goal, we adopted an objective indicator of programming expertise and recruited a population of subjects covering wide range of programming expertise. It is worth noting that the expertise indicator and behavioral/neural data obtained in this study were completely independent from each other. Because our novel laboratory task well bridged between them, we succeeded to associate programming expertise with programmers’ cortical representations in a reasonable way.

Despite the difference in research aims, a subset of brain regions specified in this study was similar to those specified by prior fMRI studies on programmers (Siegmund et al., 2014, 2017; Peitek et al., 2018a). In particular, this study associated the left IFG, MTG, IPL, SMG with programming expertise while previous studies related them with program comprehension processes. This commonality is remarkable because these results jointly suggest that both program comprehension processes and its related expertise may depend on the same set of brain regions. Providing interpretations of their potential roles in programming expertise would be beneficial for orienting future researches. First, the left IFG Tri and the left posterior MTG are frequently involved in semantic selecting and retrieving tasks (Demonet et al., 1992; Thompson-Schill et al., 1997; Simmons et al., 2005; Price, 2012). Several studies indicated that these two regions are sensitive to cognitive demands for directing semantic knowledge retrieval in a goal-oriented way (Rodd et al., 2005; Kuhl et al., 2007; Whitney et al., 2010). The involvements of the two regions in our findings may be induced by similar demands specialized for retrieval of program functional category and suggest that higher programming expertise is related to abilities of goal-oriented knowledge retrieval.

Second, many neuroscientists have shown the left IPL and SMG to be functionally related to visual word reading (Bookheimer et al., 1995; Philipose et al., 2007; Stoeckel et al., 2009) and episodic memory retrieval (Wagner et al., 2005; Vilberg and Rugg, 2008; O’Connor et al., 2010). Both cognitive functions potentially relate to the program categorization task used in our experiment. Visual word reading can be naturally engaged since source code is comprised of many English-like words and subjects may have actively recollected previously-acquired memories to compensate for insufficient clues because they had only ten seconds to categorize the given code snippet. The involvements of the left IPL and SMG in programming expertise suggest that expert programmers might possess different reading strategies and/or depend more on domain-specific memory retrieval than novices.

Other novel findings in the present study were potential involvement of the left IT, right MFG, and right IFG Tri with programming expertise. Importantly, these regions were not specified by previous studies focusing on the relationship between brain activity and program comprehension processes (Siegmund et al., 2014, 2017; Floyd et al., 2017; Peitek et al., 2018a), suggesting that the regions might be more related to programming expertise than program comprehension processes. Because the left IT is well known for the function in high-level visual processing including word recognition and categorical object representations (Chelazzi et al., 1993; Nobre et al., 1994; Kriegeskorte et al., 2008), our results may suggest that the high-level visual cortex in expert programmers could be fine-tuned by their training experience to realize faster program comprehension process. In contrast, the primary visual area showed significant decoding accuracy but no correlation to programming expertise. The evidence suggests that programming expertise could be mainly associated with high-level visual perception, although the snippets gave rise to significant activation in primary visual area.

The right MFG and IFG Tri are functionally related to stimulus-driven attention control (Corbetta et al., 2008; Japee et al., 2015). The involvement of these two regions suggests that programmers with high-level programming expertise may employ different attention strategies than less-skilled ones. Moreover, additional engagements of right hemisphere regions in experts are common across expertise studies. For example, chess experts (Bilalić et al., 2011) and abacus experts (Tanaka et al., 2002; Hanakawa et al., 2003) showed additional right hemisphere region involvements when performing their domain-specific tasks. Several fMRI studies further suggested that such activation shifts from left to right hemisphere may be related to experts’ cognitive strategy changes (Bilalić et al., 2011; Tanaka et al., 2012). Cognitive strategy changes have been observed repeatedly in comparisons between expert and novice programmers: A major characteristic is a transition from bottom-up (or textual-driven) to top-down (or goal-driven) program comprehension, which becomes feasible by experts’ domain-specific knowledge (Koenemann and Robertson, 1991; Fix et al., 1993; Von Mayrhauser and Vans, 1995). The involvement of the right MFG and IFG Tri observed in this study might be related to such cognitive strategy differences between programmers in the program categorization task.

Our results associated programming expertise with decoding accuracies of not only *category* but also *subcategory*, even though the subordinate-level categorizations were not explicitly required by the performing task. We observed that individual behavioral performances were significantly correlated with *subcategory* decoding accuracies on the left STG and SMG. These two regions are functionally related to pre-lexical and phonological processing in natural language comprehension (Demonet et al., 1992; Moore and Price, 1999; Burton et al., 2001). Interestingly, we also found a significant correlation between behavioral performances and *category* decoding accuracies on the temporal regions (left MTG and IT) associated with more semantical processing (Rodd et al., 2005; Whitney et al., 2010; Price, 2012). If these functional interpretations could be adaptable to program comprehension processes, it would be intuitive that subordinate concrete concepts (i.e. *subcategory*) of source code are processed in the left STG/SMG and more semantically abstract concepts (i.e. *category*) are represented in the left MTG/IT. This might suggest a hypothesis that an expert programmer’s brain has a hierarchical semantic processing system to obtain mental representations of source code for multiple levels of abstraction.

The results obtained via the present study were limited to a specific type of programming expertise evaluated by the expertise indicator and laboratory task used in the experiment. We particularly examined the ability to semantically categorize source code that correlated with programming expertise to win high scores in competitive programming contests. The ability to write efficient SQL programs, for example, may be an explicit indicator of another type of programming expertise but this study did not cover. Thus, our results should not be taken to imply the relationship between the neural correlates revealed here and other types of programming expertise that could not be examined by this experiment. However, it is also a fact that we cannot investigate the neural bases of programming expertise without a clear definition of expertise indicator and laboratory task that well fit the general constraints of fMRI experiments. To mitigate the potentially inevitable effects caused by this limitation, we adopted the objective indicator of programming expertise that directly reflects programmers’ actual performances and recruited a population of subjects covering a wide range of programming expertise. This study can be a baseline example for future researches to investigate the neural bases of programming expertise and other related abilities.

Our decoding framework specialized for the functional category of source code could be extended by the recent advances of decoding/encoding approaches in combination with distributed feature vectors (Diedrichsen and Kriegeskorte, 2017). Several researchers have demonstrated frameworks to decode arbitrary objects using a set of computational visual futures representing categories of target objects (Horikawa and Kamitani, 2017) and to decode perceptual experiences evoked by natural movies using word-based distributed representations (Nishida and Nishimoto, 2018). Other studies have also used word-based distributed representations to systematically map semantic selectivity across the cortex (Huth et al., 2016; Pereira et al., 2018). Meanwhile, researchers in the program analysis domain have proposed distributed representations of source code based on abstract syntax tree (AST) (Alon et al., 2019a; Zhang et al., 2019). Alon et al., for instance, have presented continuous distributed vectors representing the functionality of source code using AST and path-attention neural network (Alon et al., 2019b). The combination of recent decoding/encoding approaches and distributed representations of source code may enable us to build a computational model of program comprehension that connecting semantic features of source code to programmers’ perceptual experiences.

## 5. Conclusion

Our findings reveal a tight association between programming expertise and cortical representations of program source code in a programmer’s brain. We demonstrated that functional categories of source code can be decoded from programmer’s brain activity and the decoding accuracies on the seven regions in the frontal, parietal, and temporal cortices were significantly correlated with individual behavioral performances. The results additionally suggest that cortical representations of fine functional categories (*subcategory*) on the left SMG and STG might be associated with advanced-level programming expertise. Although research on the neural basis of programming expertise is still in its infancy, we believe that our study extends the existing human expertise literature into the domain of programming by demonstrating that top-level programmers have domain-specific cortical representations.

## Acknowledgments

We thank Takao Nakagawa and Hidetake Uwano for helpful comments on the initial study design. This work was supported by JSPS KAKENHI Grant Number JP15H05311, JP16H05857, JP16H06569, JP17H01797, JP18K18108, JP18K18141, JP18J22957, and JST ERATO Grant Number JPMJER1801.

## Author contributions

I.Y., T.K., S.Nishida, H.H., and S.Nishimoto designed the study. I.Y., T.K., and H.H. recruited subjects. I.Y., T.K., S.Nishida, and S.Nishimoto conducted experiments. I.Y., T.K., and S.Nishida analyzed data. All authors contributed interpreting results. I.Y., T.K., and S.Nishida wrote the manuscript with input from other co-authors.

