## Supplementary Information File for "Expert programmers have fine-tuned cortical representations of source code"

written by Ikutani, Kubo, Nishida, Hata, Matsumoto, Ikeda, and Nishimoto.

In this supplement, we present seven supplementary tables and five supplementary figures. Specifically, we include the following information: Supplementary Table 1 and 2 show the detailed descriptions of each category and subcategory class. Supplementary Figure 1 shows examples of Java code snippets used in the study. Supplementary Table 3 describes the statistics of the Java code snippets. Supplementary Table 4 shows the proportion of subjects who recognized each subcategory class in fMRI experiments. Supplementary Figure 2 shows the distribution of peak decoding accuracies. Supplementary Table 5 shows the locations and extents of the clusters showing significant correlations between behavioral performance and decoding accuracy. Supplementary Figure 3 shows scatter plots of peak correlation in the significant clusters. Supplementary Figure 4 and 5 show the results of category and subcategory decoding analyses using a five-voxel radius searchlight, respectively. Supplementary Table 6 and 7 show the locations and extents of the clusters showing significant correlations between behavioral performance and decoding accuracy in the five-voxel radius searchlight analyses. For further information, please contact the corresponding author ( email, ).

Supplementary Table 1. ***Description of each category***

| Name | Description provided to subjects |
|---|---|
| Math | Applying number theory processing on given inputs. |
| Search | Identifying a specific item in a list of given inputs. |
| Sort | Arranging given inputs into a certain order. |
| String | Applying a specific operation on string inputs. |

Supplementary Table 2. ***Description of each subcategory***

| Name | Description provided to subjects |
|---|---|
| Greatest common divisor | Finding the greatest common divisor of two given natural numbers. |
| Power | Calculating the powers of two given integers m and n, i.e. n-th power of m. |
| Primality test | Judging whether the given natural number is a prime number or not. |
| Binary search | A process to search for a value from the given input sequence that is equal to the target value. This process first compares the target value with the middle value of the given sequence and specify the half of the sequence that potentially contains the target value. This process iterates the comparison and sequence division until the target value is found. |
| Linear search | A process to search for a value from the given input sequence that is equal to the target value. This process examines in order from the first element of the given series. |
| Bubble sort | Arranging given inputs in a certain order by exchanging adjacent elements if they are in wrong order. |
| Insertion sort | Arranging given inputs in a certain order by iteratively picking up an element from the given input sequence and insert it to the correct location. |
| Selection sort | A process to arrange given inputs in a certain order. This process first identify the smallest value of the given inputs and exchange it with the value in the first order. Then, identify the second smallest value and exchange it with the value in the second order. This process iterates this operation until the given inputs are correctly sorted. |
| Run length encode | A process to compress the given string sequence by replacing a sequence of the same character with the character and the number of repetition. For example, the string sequence 'AAAABBBAABBBB' will be compressed as '4A3B2A4B'. |
| String sort | Sorting the given words in lexicographic order. |
| Substring search | Detecting where the given pattern appears in the given string sequence. |

**Greatest common divisor**

```
public static void main(String[] arg) {
  if (a >= b) {
    function1(a, b);
  } else {
    function1(b, a);
  }
}
static void function1(int n, int m) {
  if ((n % m) == 0) {
    System.out.println(m);
  } else if (m == 1) {
    System.out.println(1);
  } else {
    int m1 = (n % m);
    function1(m, m1);
  }
}
```

**Power**

```
public static void main(String[] args) {
  long m = scan.nextLong();
  long n = scan.nextLong();
  System.out.println(function1(m, n, 100));
}
private static long function1(long m) {
  long result = 1;
  for (long i = 1; i <= n; i++) {
    result *= m;
    if (result >= M) {
      result = result % M;
      result = function1(result, (long) n / i, 100);
      i = n - (n % i);
    }
  }
  return result;
}
```

**Primality test**

```
public static void main(String[] args) {
  int n = input.nextInt();
  int res = 0;
  for (int i = 0; i < n; ++i) {
    int x = input.nextInt();
    if (function1(x))
      ++res;
  }
}
static boolean function1(int x) {
  if (x < 2)
    return false;
  for (int i = 2; i <= Math.sqrt(x); ++i) {
    if (x % i == 0)
      return false;
  }
  return true;
}
```

**Binary search**

```
public static void main(String[] args){
    Scanner input = new Scanner(System.in);
    int n, q;
    n = input.nextInt();
    ArrayList s = new ArrayList();
    for (int i = 0; i < n; ++i) {
      int x = input.nextInt();
      if (s.size() > 0)
        continue;
      s.add(x);
    }
    s.retainAll(t);
    System.out.println(s.size());
}
```

**Linear search**

```
public static void main(String args[]) {
  int n, q, T, cnt, ans = 0;
  int[] S = new int[10001];
  for (int i = 0; i < n; i++) {
    S[i] = sc.nextInt();
  }
  for (int i = 0; i < q; i++) {
    T = sc.nextInt();
    S[n] = T;
    cnt = 0;
    while (S[cnt] != T) {
      cnt++;
    }
    if (cnt < n) {
      ans++;
    }
  }
}
```

**Bubble sort**

```
public static void main(String args[]) {
  Scanner sc = new Scanner(System.in);
  ArrayList A = new ArrayList();
  n = sc.nextInt();
  for (int i = 0; i < n; i++) {
    A.add(sc.nextInt());
  }
  for (int i = 0; i < n - 1; i++) {
    for (int j = n - 1; j > i; j--) {
      if (A.get(j - 1) > A.get(j)) {
        temp = A.get(j - 1);
        A.set(j - 1, A.get(j));
        A.set(j, temp);
        cnt++;
      }
    }
  }
```

**Insertion sort**

```
public static void main(String[] args) {
  int[] A = new int[N];
  for (int i = 1; i < N; i++) {
    int key = A[i];
    int j = i - 1;
    while (j >= 0 && A[j] > key) {
      A[j + 1] = A[j];
      j--;
    }
    A[j + 1] = key;
    System.out.print(A[0]);
    for (int k = 1; k < N; k++) {
      System.out.print(" " + A[k]);
    }
    System.out.println();
  }
}
```

**Selection sort**

```
public static void main(String[] args) {
  for (int i = 0; i < n; i++)
    a[i] = In.nextInt();
  int count = 0;
  for (int i = 0; i < n - 1; i++) {
    int minj = i;
    for (int j = i; j < n; j++) {
      if (a[j] < a[minj]) {
        minj = j;
      }
    }
    if (minj != i) {
      int tmp = a[i];
      a[i] = a[minj];
      a[minj] = tmp;
    }
  }
}
```

**Run length encode**

```
public static void main(String[] args) {
  while (stdIn.hasNext()) {
    for (int i = 0; i < t.length;) {
      if (t[i] != '@') {
        i++;
      } else {
        i += 2;
        int f = t[i - 1] - '0';
        for (int j = 0; j < f; j++) {
          System.out.print(t[i]);
        }
        i++;
      }
    }
  }
}
```

**String sort**

```
public static void main(String[] args) {
  int x, i;
  String stock;
  if (x != 0) {
    String[] data = new String[x];
    for (i = 0; i < x; i++) {
      data[i] = scan.next();
    }
    for (i = 1; i < x; i++) {
      if (data[0].compareTo(data[i]) > 0) {
        stock = data[0];
        data[0] = data[i];
        data[i] = stock;
      }
    }
  }
}
```

**Substring search**

```
public static void main(String[] args){
  String w, t;
  String[] strArray;
  int n = 0;
  w = br.readLine().toLowerCase();
  while (!(t = br.readLine()).equals("EOF")) {
    strArray = t.split(" ");
    for (int i = 0; i < strArray.length; i++) {
      if (strArray[i].toLowerCase().equals(w)) {
        n++;
      }
    }
  }
  System.out.println(n);
}
```

Supplementary Figure 1. ***Java code snippets used in the study***. 72 types of Java code snippet were used in this study. Each belonged to one subcategory and its corresponding category shown in Figure.1a. This figure shows example snippets for each subcategory class.

Supplementary Table 3. ***Statistics of Java code snippets used in the study***.

| Category | Subcategory | Number of code snippets | LOC (Mean±SD) | CPL (Mean±SD) |
|---|---|---|---|---|
| Math | Greatest common divisor | 6 | 24.8±2.2 | 59.7±19.7 |
| | Power | 6 | 25.2±3.4 | 57.2±13.9 |
| | Primality test | 6 | 26.5±1.5 | 64.8±15.8 |
| Search | Binary search | 6 | 27.2±2.4 | 67.0±17.9 |
| | Linear search | 12 | 25.6±1.2 | 57.9±22.5 |
| Sort | Bubble sort | 6 | 28.7±1.8 | 63.5±18.6 |
| | Insertion sort | 6 | 27.2±0.8 | 65.8±15.0 |
| | Selection sort | 6 | 29.1±1.0 | 54.0±13.1 |
| Sort | Run length encode | 6 | 25.0±2.2 | 52.2±14.4 |
| | String sort | 6 | 25.2±1.9 | 62.0±22.4 |
| | Substring search | 6 | 26.5±3.2 | 55.0±12.8 |

LOC: Lines of code, CPL: Characters per line.

Supplementary Table 4. ***Proportion of subjects who recognized each subcategory class in fMRI experiments***. The values were calculated based on subjects' self-reporting. Bold numbers indicate the highest value among three expertise levels. Averaged number of subcategory classes recognized by a single subject was 9.3 classes in expert; 6.7 in middle; 2.9 in novice.

| Subcategory | Expert | Middle | Novice |
|---|---|---|---|
| Greatest common divisor | **100** | 60 | 20 |
| Power | **100** | 70 | 0 |
| Primality test | **100** | 80 | 30 |
| Binary search | **100** | 70 | 30 |
| Liner search | **90** | 70 | 20 |
| Bubble sort | **100** | 70 | 50 |
| Insertion sort | 60 | **70** | 60 |
| Selection sort | **90** | 70 | 40 |
| Run Length Encode | **30** | 20 | 0 |
| String sort | **90** | 60 | 30 |
| Substring search | **70** | 30 | 10 |

Unit : [%]

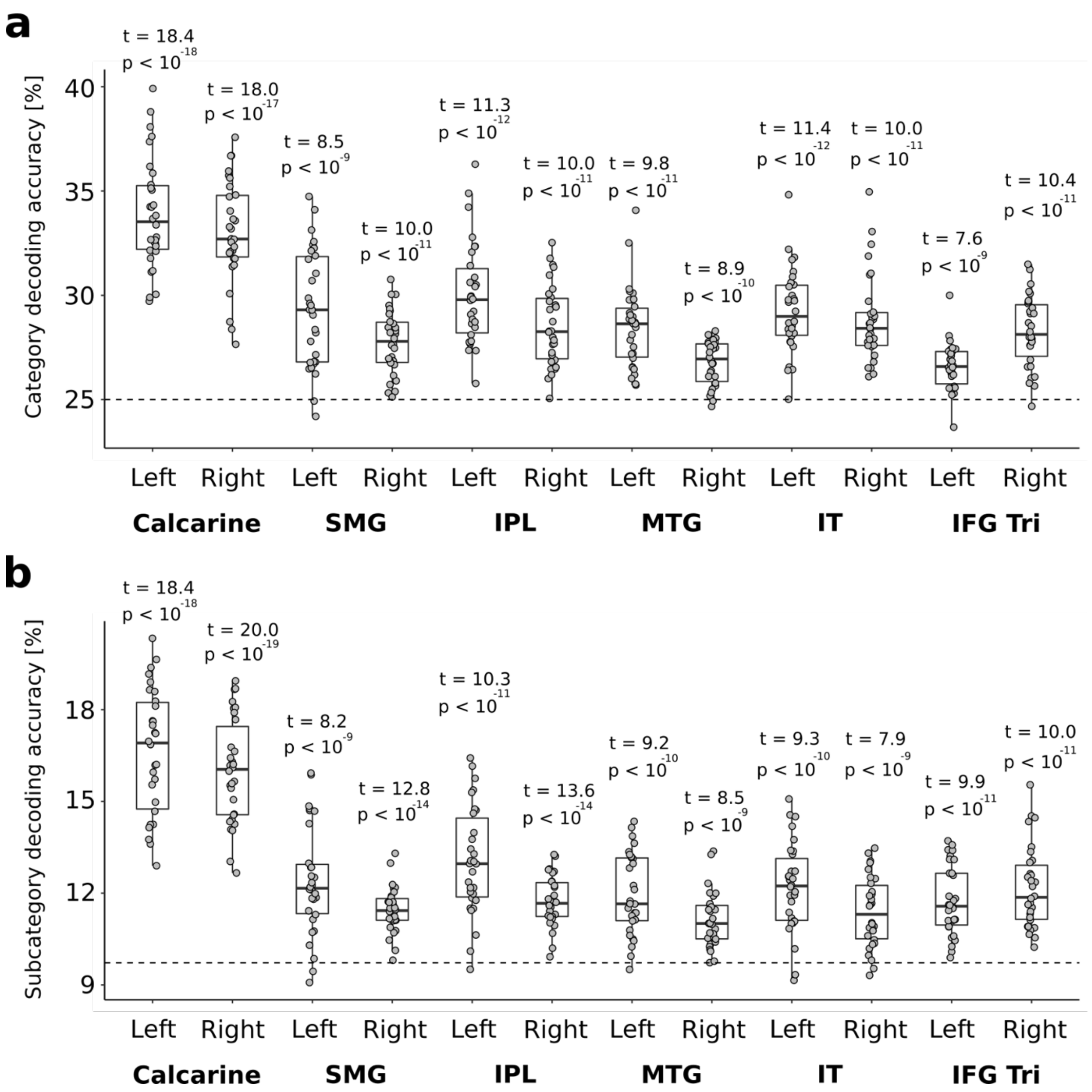


Supplementary Figure 2. ***Distribution of peak decoding accuracies***. (a) Box plots of the voxel-level peak category decoding accuracies on several brain regions. (b) Box plots of the voxel-level peak subcategory decoding accuracies. Each dot represents decoding accuracy of individual subject. The dashed line indicates chance-level accuracy; (a) 25% and (b) 9.72%. The mean decoding accuracies of all regions were significantly higher than chance determined by the second-level analyses. The regions were identified by the automated anatomical labels of Wake Forest University (WFU) PickAtlas toolbox (https://www.nitrc.org/projects/wfu_pickatlas/). Abbreviations: SMG, Supramarginal gyrus; IPL, Inferior parietal lobule; MTG, Middle temporal gyrus; IT, Inferior temporal gyrus; IFG Tri, Inferior frontal gyrus pars triangularis.

Supplementary Table 5. ***Clusters showing significant correlations between behavioral performance and decoding accuracy (voxel-level p < 0.001 and cluster-level p < 0.05, FWE-corrected)***.

| Cluster index | Extent | Corr. (***r***) | T-value | Region label | MNI coordinate | | |
|---|---|---|---|---|---|---|---|
| | | | | | x | y | z |
| 1 | 369 | 0.79 | 6.81 | R IFG (p. Triangularis) | 46 | 22 | 8 |
| 2 | 298 | 0.71 | 5.35 | L Posterior-Medial Frontal | -12 | 0 | 66 |
| 3 | 587 | 0.70 | 5.17 | R Superior Medial Gyrus | 6 | 52 | 42 |
| 4 | 649 | 0.70 | 5.16 | L Inferior Parietal Lobule | -56 | -28 | 50 |
| 5 | 428 | 0.67 | 4.84 | R Superior Frontal Gyrus | 24 | 4 | 60 |
| 6 | 346 | 0.67 | 4.79 | L IFG (p. Triangularis) | -52 | 30 | 24 |
| 7 | 347 | 0.65 | 4.50 | L Inferior Occipital Gyrus | -52 | -72 | 2 |
| | 347 | 0.64 | 4.35 | L Inferior Temporal Gyrus | -50 | -54 | 0 |

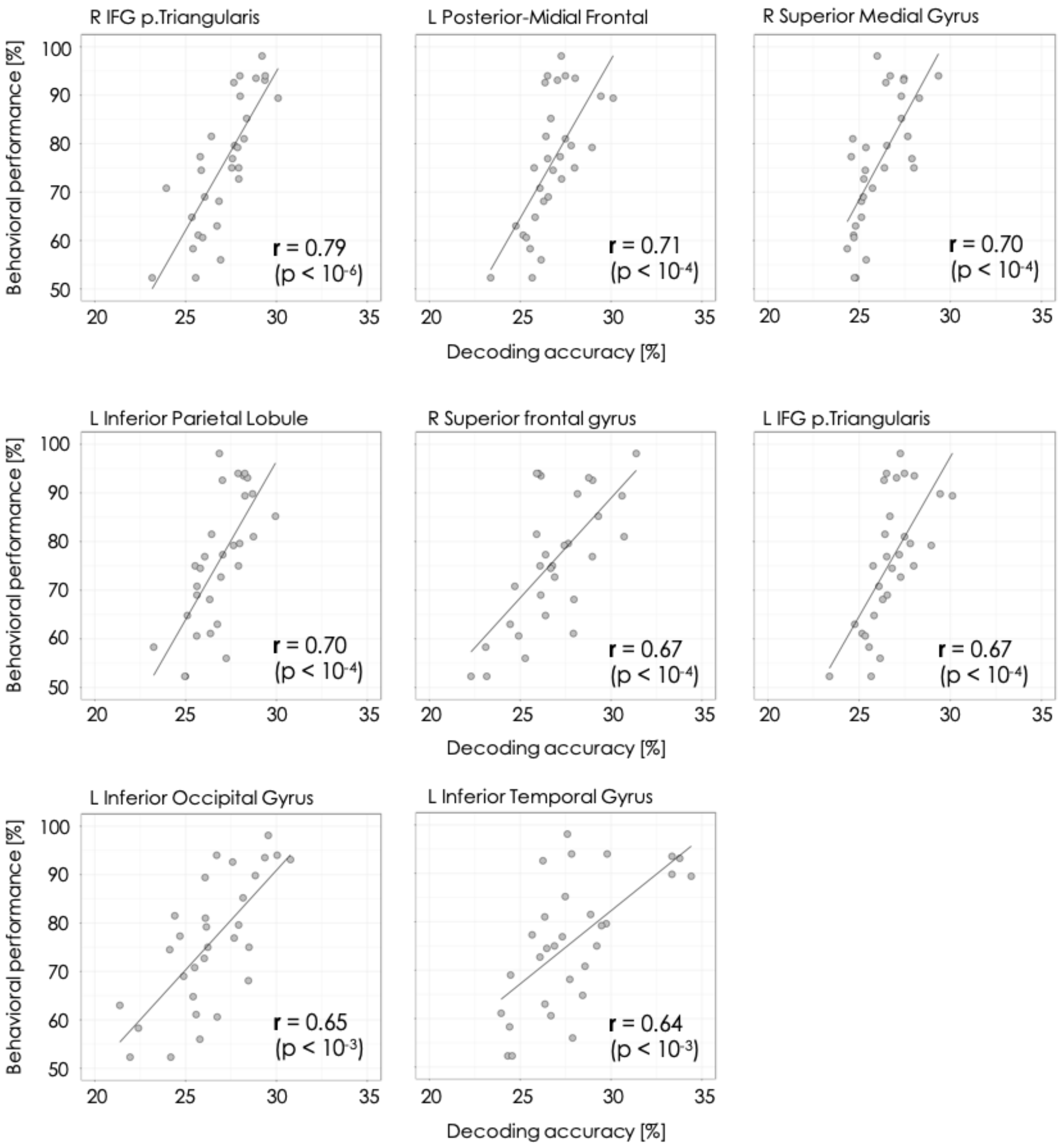


Supplementary Figure 3. ***Peak correlations of all significant clusters shown in supplementary table 5.*** Solid lines indicate fitted regression lines between decoding accuracies and behavioral performances. Correlation coefficients (***r***) and p values are shown in bottom-right of each plot.

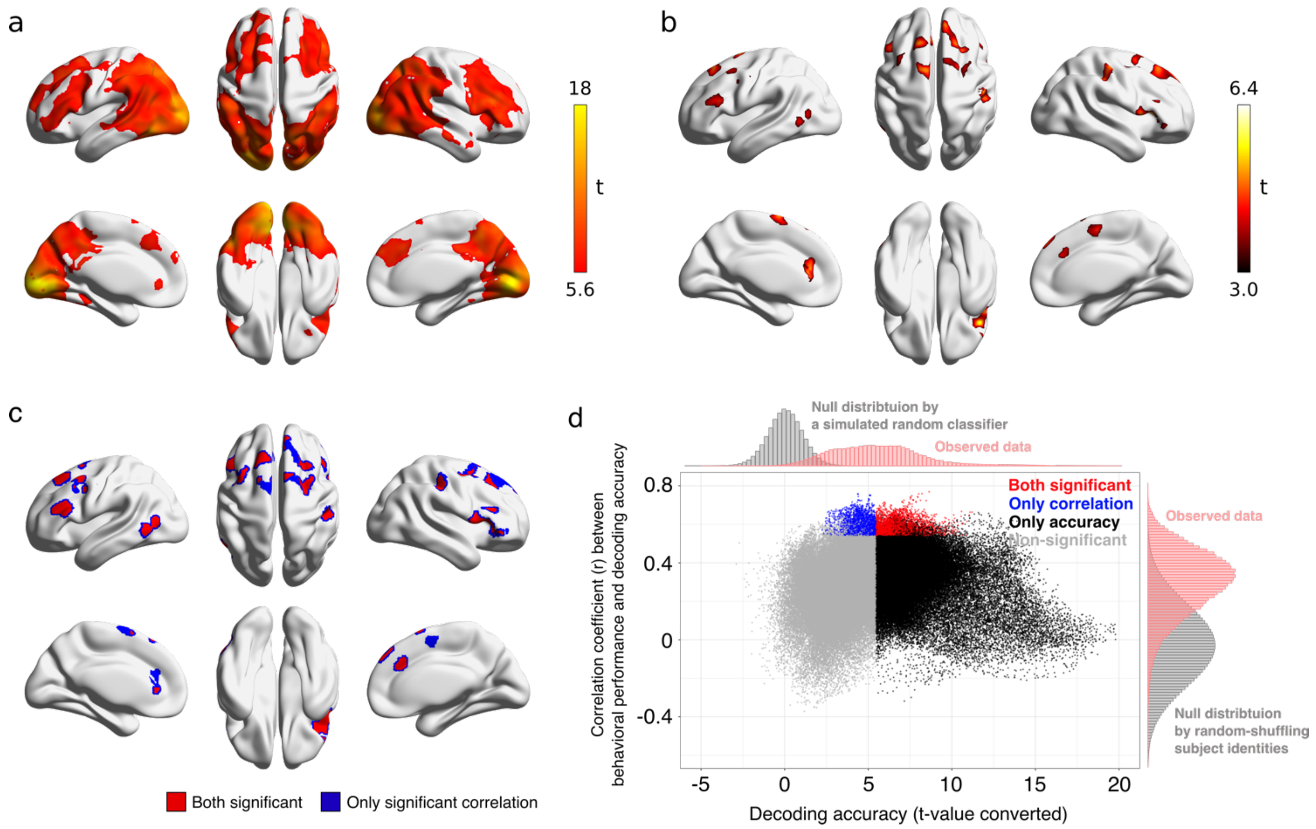


Supplementary Figure 4. ***Category decoding results using a five-voxel-radius searchlight***. (**a**) Significant searchlight locations estimated from all subject data (N = 30). Heat colored voxels denote the centers of searchlights with significant decoding accuracy (voxel-level $p < 0.05$, FWE corrected). (**b**) Locations of searchlight centers showing significant correlation between behavioral performance and decoding accuracy (voxel-level $p < 0.001$ and cluster-level $p < 0.05$, FWE-corrected). (**c**) Locations of searchlight centers that showed both significant decoding accuracy and significant correlations to behavioral performances. (**d**) Scatter plot of searchlight results. X-axis shows t-values calculated from all subjects' decoding accuracies on each searchlight locations. Y-axis indicates correlation coefficients between decoding accuracies and behavioral performances.

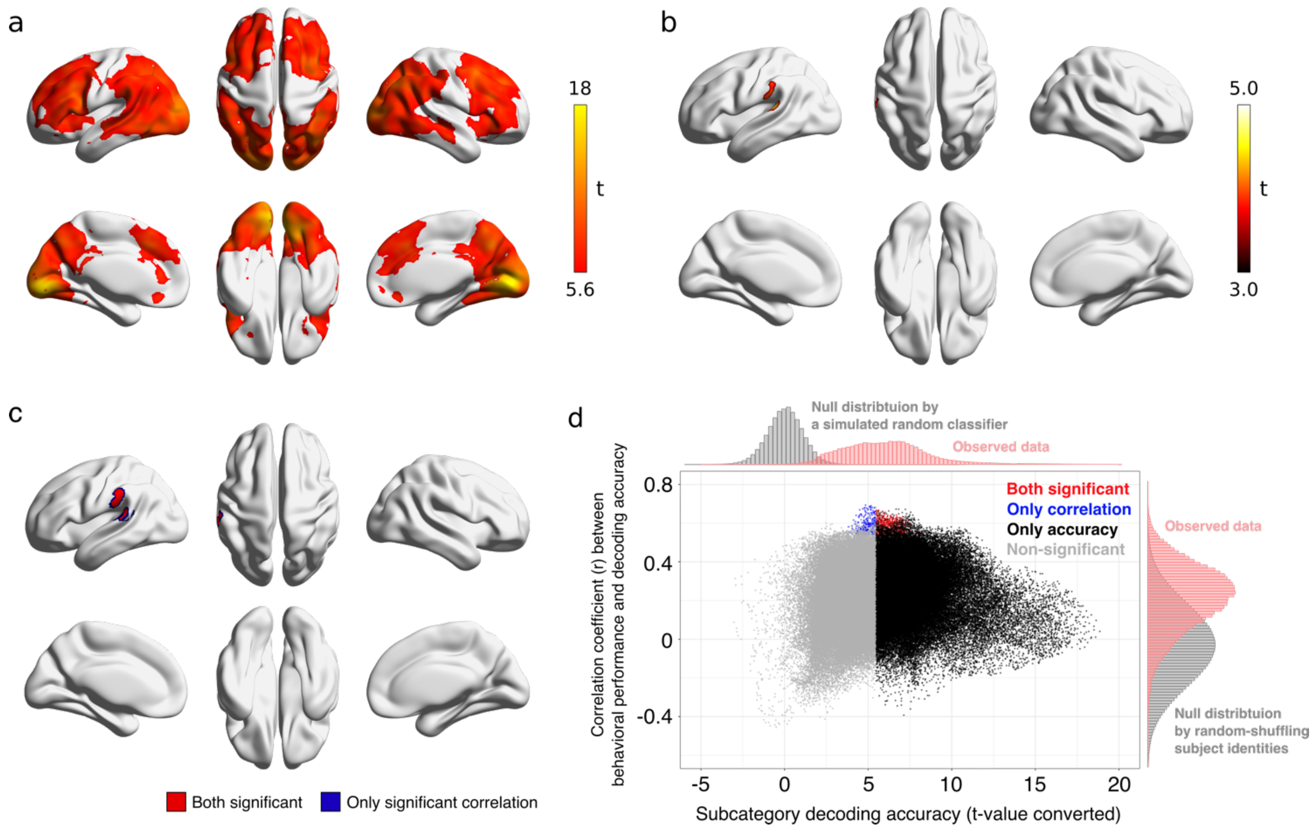


Supplementary Figure 5. ***Subcategory decoding results using a five-voxel-radius searchlight***. (**a**) Significant searchlight locations estimated from all subject data (N = 30). Heat colored voxels denote the centers of searchlights with significant subcategory decoding accuracy (voxel-level $p < 0.05$, FWE corrected). (**b**) Locations of searchlight centers showing significant correlation between behavioral performance and subcategory decoding accuracy (voxel-level $p < 0.001$ and cluster-level $p < 0.05$, FWE-corrected). (**c**) Locations of searchlight centers that showed both significant subcategory decoding accuracy and significant correlations to behavioral performances. (**d**) Scatter plot of searchlight results. X-axis shows t-values calculated from all subjects' subcategory decoding accuracies on each searchlight locations. Y-axis indicates correlation coefficients between subcategory decoding accuracies and behavioral performances.

Supplementary Table 6. ***Clusters showing significant correlations between behavioral performance and category decoding accuracy (using a five-voxel-radius searchlight; voxel-level p < 0.001 and cluster-level p < 0.05, FWE-corrected).***

| Cluster index | Extent | Corr. (***r***) | T-value | Region label | MNI coordinate | | |
|---|---|---|---|---|---|---|---|
| | | | | | x | y | z |
| 1 | 729 | 0.77 | 6.38 | ***R IFG (p. Triangularis)*** | 44 | 22 | 12 |
| 2 | 1500 | 0.76 | 6.15 | ***R Superior Medial Gyrus*** | 6 | 52 | 44 |
| 3 | 436 | 0.72 | 5.56 | R Precentral Gyrus | 48 | -20 | 52 |
| 4 | 411 | 0.70 | 5.26 | ***L Posterior-Medial Frontal*** | -12 | 2 | 66 |
| 5 | 475 | 0.70 | 5.16 | L Anterior Cingulate Cortex | -4 | 34 | 16 |
| 6 | 403 | 0.66 | 4.63 | ***L IFG (p. Triangularis)*** | -54 | 28 | 20 |
| 7 | 469 | 0.65 | 4.56 | ***L Middle Temporal Gyrus*** | -44 | -66 | 6 |
| | 469 | 0.61 | 4.10 | ***L Inferior Temporal Gyrus*** | -52 | -58 | -2 |
| 8 | 678 | 0.61 | 4.10 | ***L Middle Frontal Gyrus*** | -36 | 12 | 50 |

***Region labels in bold*** : consistent with the results using a four-voxel-radius searchlight.

Supplementary Table 7. ***Clusters showing significant correlations between behavioral performance and subcategory decoding accuracy (using a five-voxel-radius searchlight; voxel-level p < 0.001 and cluster-level p < 0.05, FWE-corrected).***

| Cluster index | Extent | Corr. (***r***) | T-value | Region label | MNI coordinate | | |
|---|---|---|---|---|---|---|---|
| | | | | | x | y | z |
| 1 | 629 | 0.69 | 5.09 | ***L Superior Temporal Gyrus*** | -56 | -32 | 14 |
| | 629 | 0.62 | 4.17 | ***L Supramarginal Gyrus*** | -64 | -32 | 38 |

***Region labels in bold*** : consistent with the results using a four-voxel-radius searchlight.
